# Rice protein models from the Nutritious Rice for the World Project

**DOI:** 10.1101/091975

**Authors:** Ling-Hong Hung, Ram Samudrala

**Author notes:** Corresponding author: Ram Samudrala. Email addresses.

## Abstract

**Background:** Many rice protein sequences are very different from the sequence of proteins with known structures. Homology modeling is not possible for many rice proteins. However, it is possible to use computational intensive *de novo* techniques to obtain protein models when the protein sequence cannot be mapped to a protein of known structure. The Nutritious Rice for the World project generated 10 billion models encompassing more than 60,000 small proteins and protein domains for the rice strains Oryza sativa and Oryza japonica.

**Findings:** Over a period of 1.5 years, the volunteers of World Community Grid supported by IBM generated 10 billion candidate structures, a task that would have taken a single CPU on the order of 10 millennia. For each protein sequence, 5 top structures were chosen using a novel clustering methodology developed for analyzing large datasets. These are provided along with the entire set of 10 billion conformers.

**Conclusions:** We anticipate that the centroid models will be of use in visualizing and determining the role of rice proteins where the function is unknown. The entire set of conformers is unique in terms of size and that they were derived from sequences that lack detectable homologs. Large sets of *de novo* conformers are rare and we anticipate that this set will be useful for benchmarking and developing new protein structure prediction methodologies.

## Data description

### Background

The function of a gene is a consequence of the protein it encodes. The three dimensional structure of a protein can be highly informative about the function and mechanism of function of the protein. Experimental determination of protein folds remains an arduous and time consuming process. However, computational methods have been developed that can predict the structure from the protein sequence. [1–3] Unfortunately, many rice proteins share little or no sequence similarity with proteins of known structure [4]. The lack of homologs meant that homology modeling was not possible for much of the rice proteome when we started this project in 2006. In the intervening time, there has been a significant increase in the number of known structures and a corresponding improvement in the coverage of homology modeling. ModBASE [5], which models whole genomes, increased the coverage of its human models from 66% in 2007 to 84% in 2013. (https://modbase.compbio.ucsf.edu/modbase-cgi/display.cgi?server=modbase&type=statistics). However, the only plant species in ModBase, *Arabdopsis,* had only 77% coverage in 2013 suggesting that even today, many plant proteins cannot by modeled using homology based methods. The lack of close homologs also means that for many genes in the rice genome, function assignment is difficult or of low confidence.[6]

In the absence of obvious homologs, it is possible to use *de novo* techniques to obtain protein models for small proteins and protein domains [1–3]. *De novo* protein modeling methods are much more computationally intensive, and less accurate than homology modeling. One method to increase the accuracy is to produce many conformers and find consensus models by clustering [7, 8].

In conjunction with World Community Grid and IBM, the Nutritious Rice for the World project (http://ram.org/compbio/protinfo/rice/) generated 10 billion *de novo* models encompassing 60,000 small proteins and protein domains for the rice strains *Oryza sativa* and *Oryza japonica.* These comprise all proteins and domains between 30-150 residues. Domains were parsed using DOMpro [9]. Volunteers downloaded the modeling software and used spare cycles to generate and upload the models to the IBM servers. These models were further refined by clustering using our local computational cluster. The project workflow is shown in figure 1.

**Figure 1.**
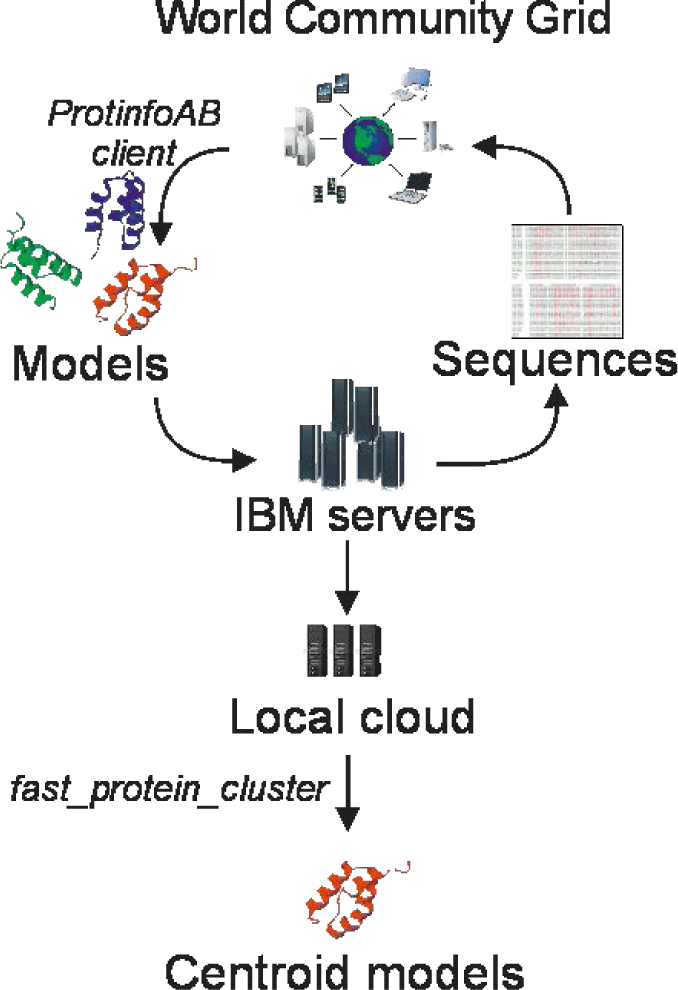
Nutritious Rice for the World workflow

This project would have taken 10 millennia for single CPU as each set of protein and protein domain conformers ranged from 100,000 to more than 550,000 in size. For the data generation we used ProtinfoAB [1, 10], which was one of the most accurate *de novo* methods in the 6^th^ Critical Assessment of protein Structure Prediction (CASP (http://www.predictioncenter.org/casp6/Casp6.html) and was one of two *de novo* servers invited to present at that meeting. We developed new methods for rapidly and accurately clustering sets of this size and have applied it to the rice conformer dataset to identify the models at the center of the 5 largest clusters. This smaller, reduced set of 300,000 represents the best predictions for the 60,000 rice proteins and rice protein domains. These folds may be useful visualizing and determining the role of rice proteins where there are no identifiable homologs and the function is unknown.

Effective protein structure prediction methods must be able to distinguish true folds from not only random coils but also from incorrect protein-like models or “decoys”, which is a much more difficult task. The set of 10 billion rice models should be useful for assessing new structure prediction methods by providing a very large background set of test models. There are very few large datasets of *de novo* conformers due to the difficulty in generating them. Also existing datasets are all based on proteins with detectable homologs which can make them problematical for the assessment of truly *de novo* techniques. The rice conformer dataset is unprecedented in the number of models per gene and the number of models derived. For example, the large Spicker [8] dataset consists of 56 sets of 10,000 to 32,000 conformers. Furthermore the rice models are derived from genes with no known homologs and unlike most other *de novo* methods, ProtinfoAB does not copy coordinates or angles from known structures. One use for these sets is the development of high resolution energy functions for protein folding. It is much more difficult for these functions to be able to distinguish true folds from compact protein-like conformers than from random coils. Our set of conformers is an even better test, in that the conformers do not contain any existing substructures from the known protein. This challenging dataset will be very useful for the development of new *de novo* structure prediction techniques.

### Data Generation

ProtinfoAB was used for generating conformers. A Monte Carlo simulated annealing (MCSA) search is conducted to find a conformation that minimizes a simple 3-term statistical energy function is minimized. The initial search is biased to dihedral angles that are found in known protein structures but no coordinates from known structures are actually copied. Once a minimum has been found, a second local search further minimizes the energy of the conformer. This is a very fast implementation (about 1 minute on a Pentium 4 per conformer) with minimal memory requirements (25 MBytes). The minimal demands allowed the software to run unobtrusively even when the donated computer time was on older hardware.

World Community Grid volunteers downloaded the ProtinfoAB app onto their computers and generated conformers for a given rice protein sequence. These were then transmitted to IBM servers where they were collected and aggregated into compressed tarballs that were sent to our servers. 150,000 to 550,000 structures were generated per sequence. The structures were stored as compressed dihedral angles rather than Cartesian coordinates to save space. As a result, the entire dataset is less than 6 TB in size.

After World Community Grid produced the set of conformers we used our local cluster to find the structures that were least dissimilar to the other structures generated from the same sequence. After benchmarking several strategies [11, 7, 12] we found that by complete linkage hierarchical clustering Root Mean Square Deviation (RMSD) after optimal superposition as the metric of structural similarity produced the best results [7]. Due to the size of the clusters, we developed new parallel, and vectorized software to implement this strategy on multi-core servers and Graphics Processing Units (GPUs) [7] (https://github.com/lhhunghimself/fast protein cluster). The centroids from the 5 largest clusters (out of 10 clusters total) were kept as the best representative structures of the entire ensemble and are made available through a simple webserver.

### Conformer sets

The raw database consists of compressed (bzip) tarballs of the dihedral angles of the conformers. This is much more efficient than storing the atomic coordinates. The database of conformers takes less than 6 TB in size. The number of best centroid structures is considerably smaller and these are stored as compressed pdb files totaling 2 GB in size. Because MD5 hashes are almost always unique, we used the MD5 hash of the sequence to name the files themselves. A sqlite database provides a mapping between the MD5 naming system, the sequence from which the MD5 hash was derived and the gene from which the sequence was derived.

This database is available through http://protinfo.org/rice/data/. Individual genes can be queried by entering the rice gene name or a protein subsequence. The unique MD5 identifiers are displayed and the user is provided a link for downloading the centroid structures matching the query. If there is more than one domain matching the gene, all matching domains are displayed. The entire database of centroid structures is also available for download on the website. Arrangements can also be made to obtain the entire database of conformers from the corresponding author In addition we provide metadata (gene locus and sequence) to allow the user to find the matching filename from the locus or sequence. In the time that we started this work, the annotation of the rice genome has evolved and some of the sequences and identified loci from that time may not correspond to what are now considered to be the real genes and proteins. Therefore, we have also provide a fasta file and a BLAST formatted database to allow sequence based queries against the conformers independent of the gene identification and nomenclature.

### Concluding remarks

The Nutritious Rice for the World project has produced a set of rice protein models that is unique in scope and size. Protein models on a genomic scale are possible but largely rely on homology modeling. Plant proteomes are difficult to model by homology modeling due to the divergence of the protein sequences. Modeling by *de novo* methods is difficult due to the increased computational needs. World Community Grid has provided us the resources to produce a set of models derived sequences that lack detectable homologs. We anticipate that the smaller set of centroid models will be of use in visualizing and determining the role of rice proteins where the function is unknown. We anticipate that the entire set of 10 billion conformers will be very useful for benchmarking and developing new protein structure prediction methodologies.

**Abbreviations used**
MCSA: Monte Carlo Simulated Annealing
CASP: Critical Assessment of Protein Structure Prediction
GPU: Graphics Processing Unit
CPU: Central Processing Unit
IBM: International Business Machines
RMSD: Root Mean Square Deviation after optimal superposition

## Authors’ contributions

LHH conceived the experiments, wrote the software and webserver and drafted the manuscript. RS conceived the project, and help draft the manuscript.

## Acknowledgements

We would like to thank the Michal Guerquin, Mike Shannon, and Haychoi Taing for setting up and maintaining the infrastructure to receive, store and analyze the data. We would like to thank the IBM team for their help and patience in setting up and moving this project forward. Finally we would like to thank all the volunteers who donated their computer time for the generation of structures and participated in the forums. Without their efforts, none of this would have been possible. This work was supported by National Institutes of Health Pioneer Award DP1LM011509 to R.S.

